# A pangenome framework uncovers the role of deletions in repeated evolution of cave-derived traits

**DOI:** 10.1101/2025.11.25.690530

**Authors:** Emma Y. Roback, X Maggs, Edward S. Ricemeyer, Adam Warlen, Rachel A. Carroll, Christine G. Elsik, Alex C. Keene, Nicolas Rohner, Suzanne E. McGaugh, Wesley C. Warren

## Abstract

Structural variants (SVs) are increasingly recognized as key contributors to adaptive evolution, yet they remain underexplored compared to single-nucleotide variation. To understand how large-scale genomic changes shape repeated evolution, we leveraged multiple levels of sequence data across the powerful evolutionary model system of the Mexican tetra fish (*Astyanax mexicanus*). Cavefish populations have repeatedly evolved similar and dramatic phenotypes compared to surface fish such as eye loss, pigmentation loss, and sleep loss, suggesting a potential role for loss-of-function variants in cave adaptation. We constructed one of the first pangenome graphs from a naturally evolving vertebrate, enabling comprehensive discovery of SVs among 120 fish from 11 populations. We identified over 2,400 high-confidence cave-specific deletions that are enriched in biological pathways involved in vision, metabolism, and behavior and cluster non-randomly in quantitative trait loci linked to cavefish traits. Strikingly, 67 genes harbor unique deletions between independent cavefish lineages. These reused genes show strong evidence of population-specific selection (99% are under selection, compared to 8–15% in genes lacking SVs), indicating that deletions rose in frequency through repeated positive selection rather than drift. Together, these results reveal that recurrent deletion events have repeatedly contributed to the evolution of cave-adapted phenotypes and highlight deletions as underexplored contributors of adaptive evolution in a system characterized by trait loss.

## INTRODUCTION

A central goal in biology is to understand how genetic variation shapes phenotypic diversity, particularly in the context of adaptation. Single nucleotide polymorphisms (SNPs) are abundant, readily detected with existing technology, and have been extensively studied for their association with traits that have evolutionary and ecological relevance (Saunders et al. 2006; Williams and Oleksiak 2011). Yet, SNPs represent only a fraction of the genetic variation that can influence phenotype (Huddleston and Eichler 2016). Much of the remaining, and historically overlooked, variation arises from large-scale genomic alterations collectively denoted as structural variation.

Structural variants (SVs) are generally defined as sequence variants ≥ 50bp in size (Sudmant et al. 2015). These variants include inversions, translocations, duplications or insertion of novel sequence, and deletions. Due to their large size and abundance in the genome, SVs account for substantially more sequence variation than SNPs (Huddleston et al. 2017; Catanach et al. 2019; Hämälä et al. 2021). Consequently, SVs may be particularly influential in shaping traits under strong selection pressure, as seen in hypoxic conditions at high elevation (Liang et al. 2024), and in cases of strong artificial selection in domesticated animals (Cumer et al. 2021; Andersson and Purugganan 2022; Liu et al. 2023). Convergent SVs have been identified in sheep and goats subject to selection for similar agricultural traits (Yang et al. 2024) and play a key role in rapid and repeated adaptation in ragweed (Battlay et al. 2025). Even loss-of-function SVs like deletions, which are generally presumed to be deleterious to fitness, have the potential to generate evolutionary novelty (Dwivedi et al. 2019; Murray 2020) and contribute to adaptation (Albalat and Cañestro 2016; Sharma et al. 2018; Xu and Guo 2020; Monroe et al. 2021).

The large size and complexity of SVs, combined with the limitations of short-read sequencing and reduced detection sensitivity, have hindered accurate genome-wide discovery, genotyping, and analysis of SVs among populations (Huddleston and Eichler 2016; Sedlazeck et al. 2018; Mahmoud et al. 2019; Zhao et al. 2021). Fortunately, recent advances in pangenome graph methods now provide a more comprehensive framework for identifying and characterizing segregating SVs. First conceptualized by (Tettelin et al. 2005), pangenomes are an alignment of multiple contiguous whole genome assemblies that can capture more of the genomic diversity that exists within a species (Bayer et al. 2020; Eizenga et al. 2020). In contrast to mapping to a linear reference, pangenomes enable genotyping of SVs from short-read data (Hickey et al. 2020) and reduce false negatives in the discovery of polymorphic SV alleles (Rice et al. 2023). For these reasons, pangenomes are the most informative approach to discover SVs that may affect phenotypes of interest. While pangenomes have been generated for numerous agriculturally relevant species (Hirsch et al. 2014; Golicz et al. 2016; Montenegro et al. 2017; Zhou et al. 2022; Jiang et al. 2023; Li et al. 2023; Rice et al. 2023), this approach has been used for only a few naturally evolving populations (Ruggieri et al. 2022; Secomandi et al. 2023; Fang and Edwards 2024; Edwards et al. 2025; Quah et al. 2025). Existing pangenomes of wild populations have revealed links between SVs and adaptation, including nervous system development in *Heliconi<u>u</u>s* butterflies (Ruggieri et al. 2022), and disease resistance in the house finch (*Haemorhous mexicanus*; Fang and Edwards 2024), illustrating the power of this approach to uncover important and previously obscured genetic variation.

We generated a pangenome graph for a unique evolutionary model system, the Mexican tetra (*Astyanax mexicanus*). The Mexican tetra is a species of teleost fish that has independently colonized cave environments at least twice in its evolutionary history. As a result, Mexican tetra have two distinct lineages (Lineage 1 and Lineage 2) each comprising both surface-morph and cave-morph populations (Herman et al. 2018; Garduño-Sánchez et al. 2023; Moran et al. 2023). Cave populations likely diverged less than 200 thousand generations ago from surface populations but continue to undergo gene flow with surface populations and both intra- and inter-lineage cave populations (Herman et al. 2018).

Subterranean habitats, defined by perpetual darkness, low oxygen, and limited food availability, have repeatedly produced organisms with reduced eyes and pigmentation, as well as expansion of non-visual senses (Schilthuizen et al. 2005; Niemiller et al. 2008; Klaus et al. 2013; Bloom et al. 2014; Yang et al. 2016; Behrmann-Godel et al. 2017). Like other subterranean organisms, Mexican cavefish consistently show loss of vision (Yamamoto et al. 2004; Krishnan and Rohner 2017) and pigment (Protas et al. 2006; Gross et al. 2009). Beyond these hallmark cave traits, Mexican cavefish have also lost the capacity to regenerate normal heart tissue after injury, an ability seen in surface fish and ancestral teleosts (Stockdale et al. 2018; Cutie and Huang 2021). In response to nutrient limitation, Mexican cavefish have altered metabolism (Riddle et al. 2018; Xiong et al. 2018; Medley et al. 2022; Xiong et al. 2022) and exhibit behavioral changes including loss of social schooling (Kowalko et al. 2013), reduced aggression (Elipot et al. 2013), and hyperphagia (Aspiras et al. 2015). Although caves differ somewhat in their ecology (Mitchell et al. 1977; Elliott 2018), adaptation to a resource limited, perpetually dark cave environment has proceeded in a similar manner at the phenotypic level across cavefish populations (Morris 2003; Hernández-Lozano et al. 2024).

The repeated evolution of similar phenotypes in Mexican cavefish has long intrigued evolutionary biologists, prompting efforts to uncover the underlying genetic basis of cave-derived traits. To this end, previous research has identified a handful of genes associated with cavefish phenotypes (reviewed in Ponnimbaduge Perera et al. 2023) and a few SNPs that are thought to contribute to these traits. For example, missense mutations in *mc4r* and *insra* affect appetite and insulin signaling, respectively (Aspiras et al. 2015; Riddle et al. 2018), and a premature termination codon in *pde6c* impairs vision (Roback et al. 2025). These discoveries highlight the potential relevance of a few SNPs, but there remains a substantial gap in our understanding of the genetic basis of cave-derived traits. We address this gap through a systematic exploration of structural variation in 120 Mexican tetra genomes and identify convergence in structural variation at both the level of the gene and biological pathway.

We present the first pangenome graph of genome-wide changes associated with cave colonization, discover substantial amounts of structural variation, and explore the roles of genomic biases and selection in shaping the distribution of SVs. We show that 1) SVs occur non-randomly within the genome and are more common in long genes that have greater mutational opportunity and are more likely to be reused during independent evolutionary events; 2) cave-specific deletions are enriched within QTL for cave traits and affect genes and pathways with established links to cavefish biology; these are likely to contribute to cave-derived phenotypes; 3) unique deletions have arisen repeatedly and persisted in the same genes after cave colonization, and 99% of these genes bear evidence of selection in caves, suggesting that this pattern is not simply the result of relaxed constraint, but reflects repeated, lineage-specific selection on parallel genetic changes.

## RESULTS

### Overview of study design

Our sequencing and filtering strategy enabled us to examine SVs that likely arose after cave populations diverged from surface populations and compare SVs across independently derived cave lineages. Overall, using a total of 120 *Astyanax mexicanus* genomes from four surface and seven cave populations, we identified structural insertions and deletions originating from two independent evolutionary events (Figure 1A-D). These populations were selected to provide a robust sampling of cave and surface fish in each lineage (n= 56 for Lineage 1, n = 64 for Lineage 2), which allows for inter- and intra- lineage comparisons of structural variation.

**Figure 1.**
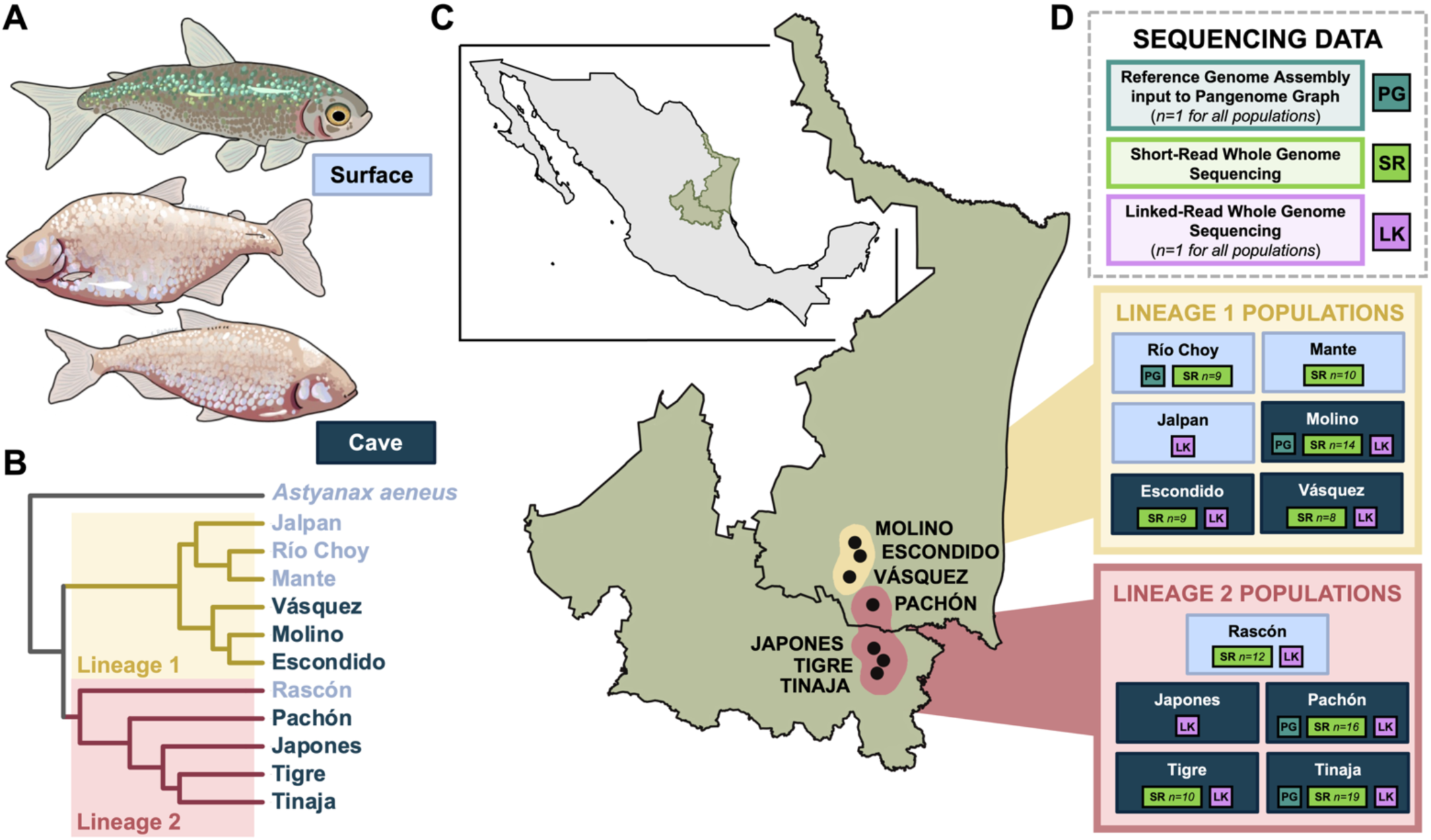
**A)** Representation of surface and cave-morph Mexican tetra fish **B)** Phylogenetic relationships of populations used in the present study (subset from full phylogeny constructed by Moran et al. 2023). Populations belong to two distinct lineages, with cavefish evolving in each lineage. **C)** Map of cave locations, San Luis Potosí and Tamaulipas, Mexico. **D)** Sequencing datasets and sample sizes for each population.

To identify SVs, we first generated a pangenome graph from four reference genomes (Imarazene et al. 2021; Warren et al. 2024) and identified insertion and deletion variants within the graph (Figure 2A). We refer to these variants as pangenome SVs and characterize them across the genome. To enable further analyses, we obtained genotype information for variants across populations by mapping short-read whole genome sequencing (WGS) data to the pangenome graph (Figure 2B). We refer to this dataset as “PGsr” (pangenome + short read WGS). In addition, we completed separate SV calling on linked-read WGS data (Figure 2B). Given that many cavefish phenotypes are losses of ancestral traits seen in surface fish (e.g., loss of eyes, pigment, and social behavior) and that SV deletions are predicted to have a high functional impact, we focused all further analyses on the study of deletions. To identify variants phenotypically relevant to cavefish, we isolated deletions that were statistically associated with cave-morph fish in each lineage. From these deletions we identified those that were consistently identified across different SV calling methods, not sampled in surface populations, and high frequency (deletion allele frequency ≥ 0.8) in cave populations (Figure 2C). We refer to this final dataset of most-filtered deletions as “focal deletions”. In some tests below, we compare genes with focal deletions to “control genes,” which contain no SV insertions or deletions within the gene structure in any population in the pangenome. For full study details, please see materials and methods.

**Figure 2.**
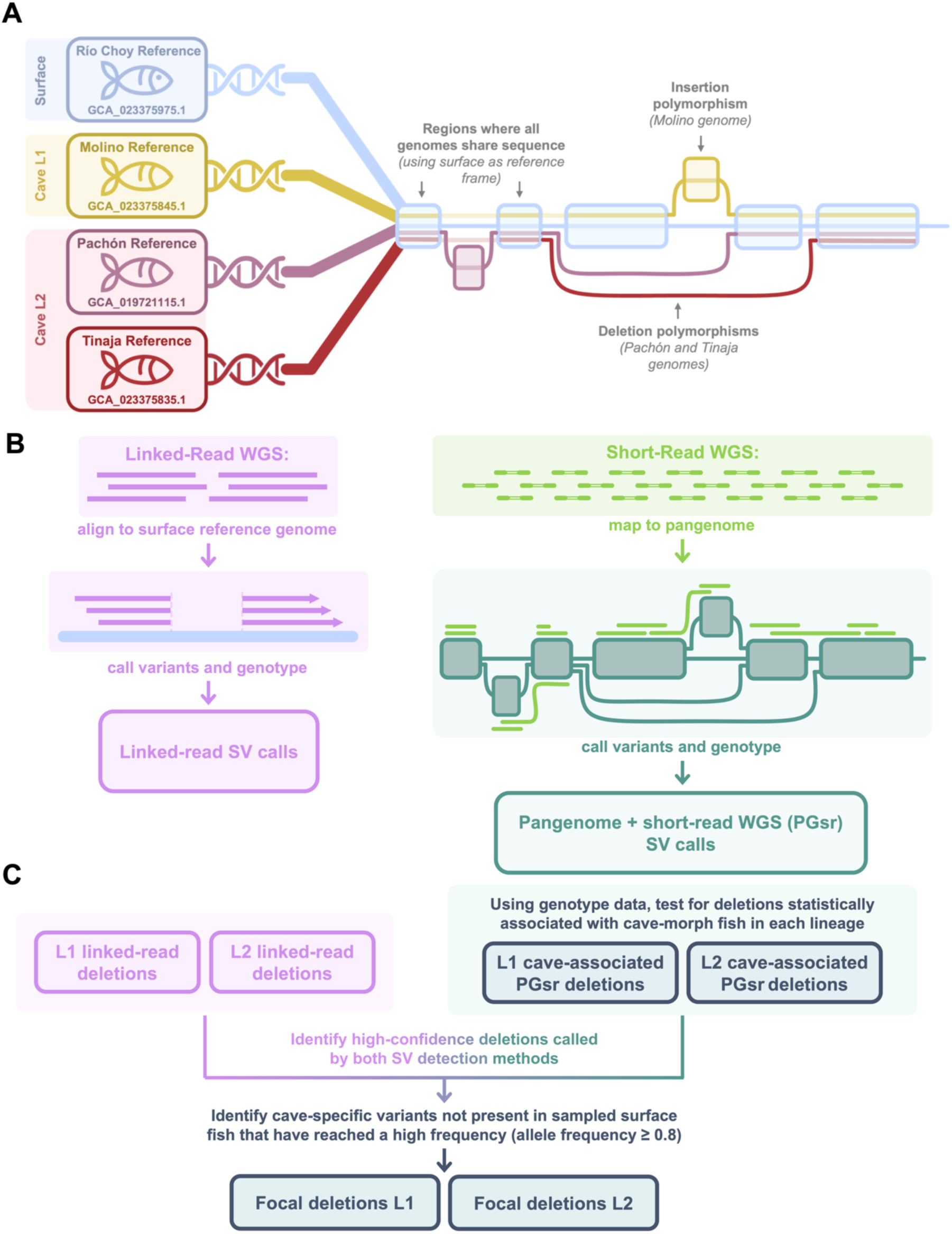
**A)** Graphical representation of pangenome graph construction using surface fish genome as a reference frame, and examples of insertion and deletions represented in the pangenome graph. **B)** SV calling methods from linked-read sequencing (left) and mapping short-read sequencing to pangenome graph (right). **C)** Integration of SV calls from different calling methods and filtering of SV deletion variants to generate a set of focal deletions used for downstream analyses.

### Pangenome graph construction

Our Mexican tetra pangenome graph, generated from reference genomes of one surface and three cave morph individuals, consists of 52,571,860 nodes (sections of DNA sequence), and 72,140,066 edges (connections between nodes indicating that the given sequences are adjacent to one another in at least one genome). During graph construction, addition of the three cave-morph genomes to the pangenome graph contributed approximately 400 Mbp of non-surface sequence, attributable to 176 Mbp in Tinaja, 131 Mbp in Molino, and 94 Mbp in Pachón. We did not reach a plateau in new non-reference sequence nodes, indicating that inclusion of additional genomes would continue to grow the pangenome graph. The pangenome graph used the Mexican tetra Río Choy surface fish reference (Warren et al. 2024) as a reference frame; thus, all results from the pangenome graph describe variation occurring in the three cave references relative to the Río Choy surface reference.

### Characteristics of pangenome insertions and deletions

From the pangenome graph, we identified variant sites: positions in the surface reference at which the cave reference genomes differ. For top-level indel variants (i.e., primary variants that are not nested within other variants, LV=0) on chromosomes specifically, we identified a total of 1,100,450 sites with one-base pair indels, 1,359,534 sites with small insertions (<50 bp), 1,489,289 sites with small deletions (<50 bp), 236,786 sites with SV insertions (sequence gains 50bp-100,000bp), 216,339 sites with SV deletions (sequence losses 50bp-100,000bp). We note that some sites affected by SVs are multiallelic, with unique SV alleles between caves. Across SV insertion sites, we found 338,631 unique insertion alleles and across SV deletion sites we found 255,331 unique alleles (Figure 3A). SV alleles were more often shared within lineages than between lineages (Lineage 1 = Molino, Lineage 2 = Pachón & Tinaja). Alleles for structural insertions amount to an average of 138 Mbp per cave morph, while alleles for structural deletions amount to an average of 106 Mbp per cave morph (Figure 3B). We find more total bp affected by SVs in the Lineage 2 populations, Pachón and Tinaja, likely because the surface reference is from a Río Choy individual which belongs to Lineage 1 and is therefore more closely related to the Molino cave population. The majority (80.31%) of SV insertions and deletions we identified in the pangenome are ≤ 1 kb in length (Supplementary Figure 1A).

**Figure 3.**
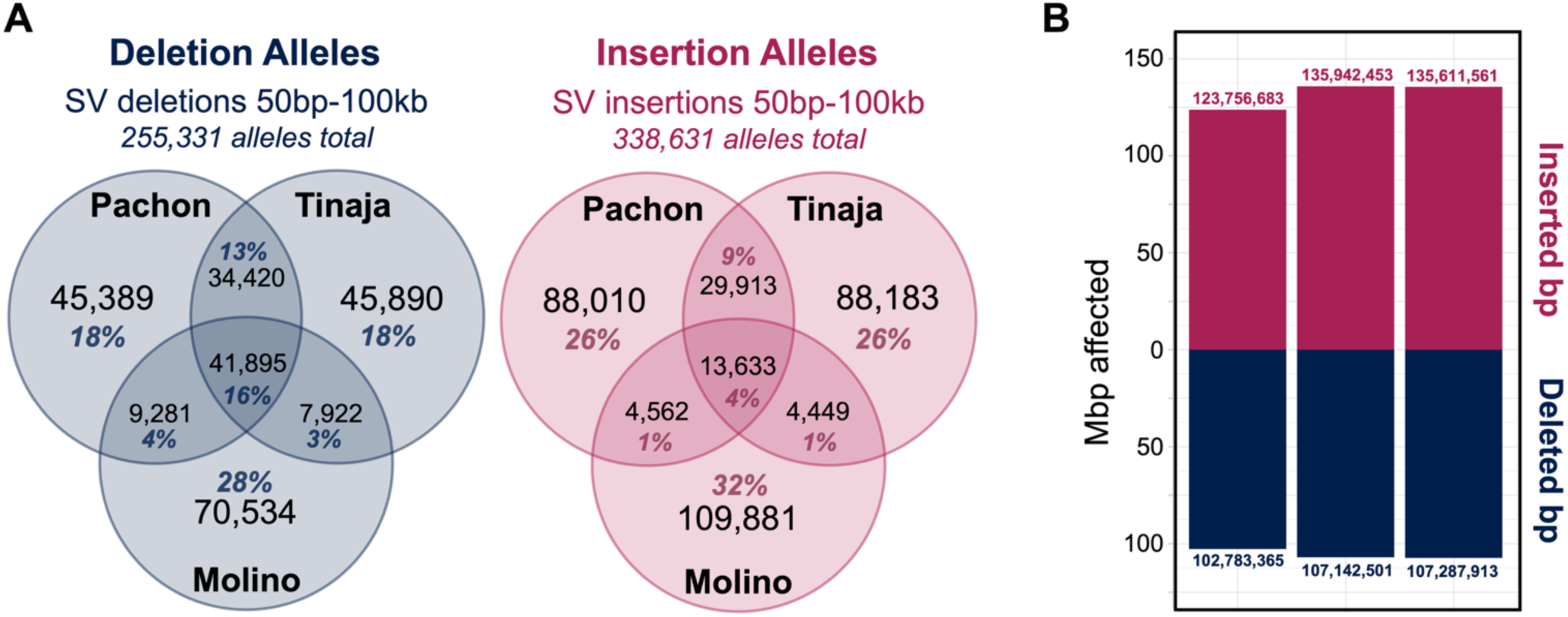
**A)** Comparison of pangenome SV deletion alleles (left) and SV insertion alleles (right) by presence in each cave population (Lineage 1 = Molino, Lineage 2 = Pachón & Tinaja). **B)** Total number of base pairs affected by pangenome SV insertions (positive values) and SV deletions (negative values) in each population.

Across the 26,736 Mexican tetra protein coding genes, 18,016 (67%) have SV deletions and 17,715 (66%) have SV insertions in Molino, Pachón, or Tinaja within intron, exon, or UTR regions. Longer genes offer more mutational targets and are therefore more likely to contain variation (Lopes et al. 2021), a trend that is well-supported in Mexican tetra (Moran et al. 2023; Roback et al. 2025). Likewise, we found that genes that contain no SV insertions or deletions are shortest in total base pairs (median = 4,847bp), followed by genes with SV insertions (median = 6,538bp), then genes with SV deletions (median = 7,177bp), and finally genes with both SV insertions and deletions (median = 23,880bp) (all groups different from one another through pairwise comparisons using Wilcoxon rank sum test with continuity correction, all p < 0.01, average Cohen’s d across all comparisons = 0.33, Table S1-A). For genes that contain SV deletions, we find a bimodal distribution where most genes have 25% or less of their total base pairs deleted, as well as a notable group of genes that are completely deleted (Supplementary Figure 1B).

Overall, 50.85% of all SV deletions and 41.55% of all SV insertions affect introns. In contrast, relatively few SV insertions and deletions occur in exons. Across the three cave morphs in the pangenome, SV deletions affect exons of 2,397 protein coding genes and SV insertions affect exons of 1,332 protein coding genes. We find that SV insertions and deletions overlap CDSs less frequently than random (permutation test p < 0.005 for both insertions and deletions, Table S1-B; Figure 4A) and overlap introns more frequently than random (p = 0.04 for deletions and p < 0.005 for insertions, Table S1-B; Figure 4B). In intergenic regions (i.e., regions of the genome not within the gene structure), there are more insertions and fewer deletions than expected through random accumulation (p < 0.005 for both insertions and deletions, Table S1-B). These results suggest that negative selection is likely removing SVs from coding regions, in line with findings in other animals (Liang et al. 2024).

**Figure 4.**
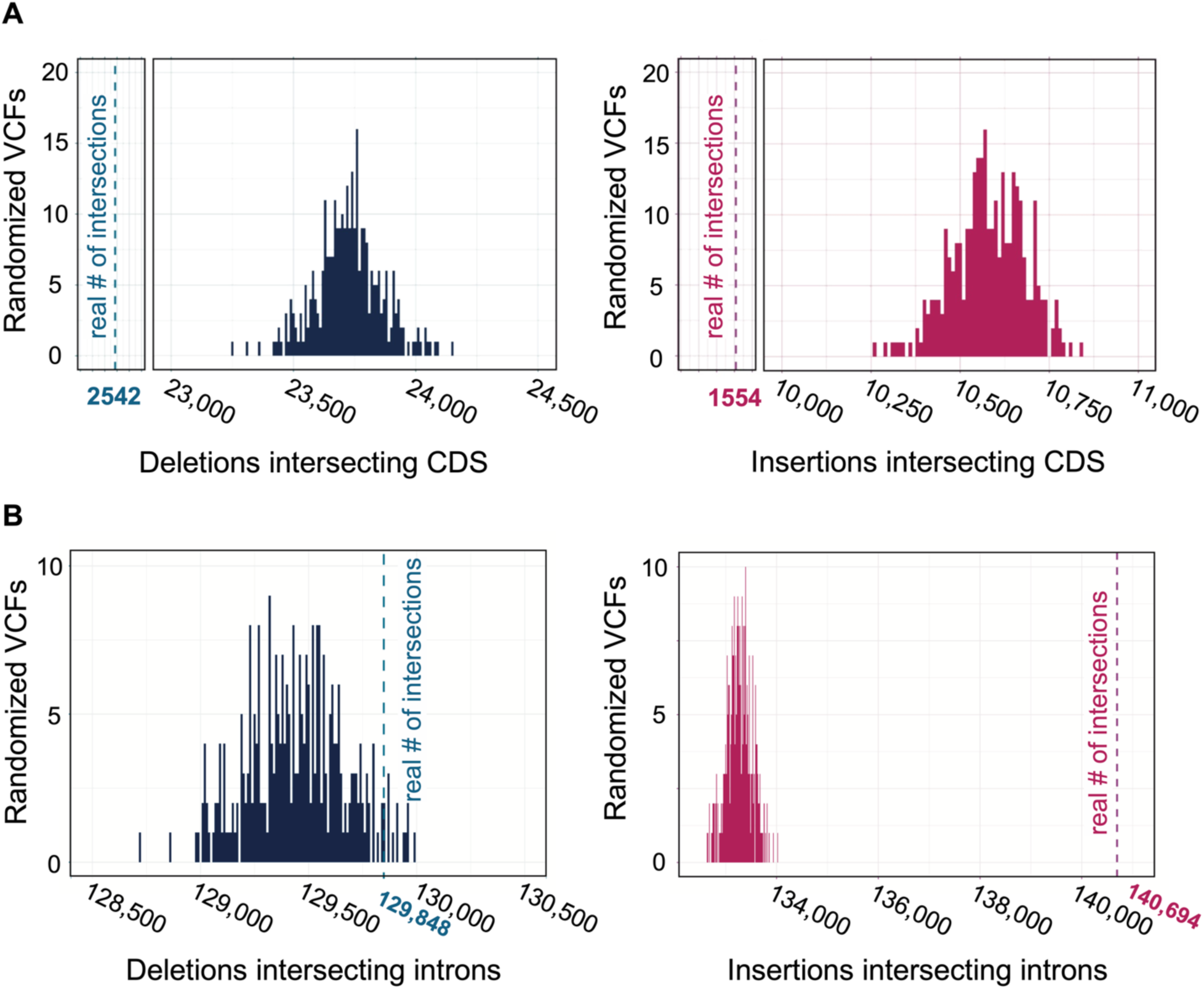
Results of permutation test that places identified SVs into the genome at random and determines how frequently the variant overlaps regions of interest. The histograms illustrate the number of intersections between the SV and the given region in 300 runs of the random permutation test and the dotted line indicates the observed number of intersections. **A)** Intersections of deletions (left) and insertions (right) with coding sequence (CDS) regions. **B)** Intersections of deletions (left) and insertions (right) with introns.

Because transposable elements can amplify and insert multiple times, they are inherently repetitive and abundant among insertion events. True to these expectations, SV insertions found in cavefish relative to the Río Choy surface reference are 78.6% repetitive on average across Molino, Pachón, and Tinaja (Table S1-C). Of this repetitive content, the largest contributors are retroelements (30.7% on average, including 21.3% from LTR elements) and DNA transposons (19.3%).

Finally, we predicted functional impacts of variants in the pangenome as characterized by SnpEff (Cingolani et al. 2012). Notably, SV deletions represent 30.92% of all high-impact variants, and SV insertions account for 10.65% (Supplementary Figure 1C; Table S1-D).

### Identification of SV deletions from linked-read sequencing

We also used linked-read sequencing to identify SV deletions in individuals from each of two surface populations and seven caves relative to the Río Choy surface reference genome (Table 1; Table S2). We obtained fewer deletion calls in Molino, Pachón, and Tinaja through the linked-reads compared to the pangenome graph (approximately 1.7X, more deletions were identified in the pangenome graph in Molino, and 1.5X more in Pachón and Tinaja). This highlights the sensitivity of SV detection provided by the pangenome graph method compared to use of a linear reference, as also demonstrated by (Rice et al. 2023). From the linked-reads we identified the fewest deletions in the Jalpan surface individual (75,975 deletions) which is the most closely related population to the Río Choy reference (both Lineage 1 surface populations). We found the most deletions in the Rascón surface individual (94,619 deletions) which represents the divergence between Lineage 1 and Lineage 2, the deepest divergence time in *A. mexicanus* (Herman et al. 2018). Additionally, this highlights the high degree of structural diversity within surface morphs from different lineages.

**Table 1.** Number of SV deletions called in each individual from linked-read sequencing dataset after filtering for variants passing all Longranger quality filters, and total number of surface reference base pairs deleted by total SV deletions.

|  | Lineage 1 |  |  |  | Lineage 2 |  |  |  |  |
| --- | --- | --- | --- | --- | --- | --- | --- | --- | --- |
|  | Escondido | *cave*<br>Molino | Vásquez | *surface*<br>Jalpan | Pachón | *cave*<br>Tigre | Tinaja | Japones | *surface*<br>Rascón |
| # Deletions (50bp-100kb) | 80,149 | 76,886 | 82,390 | 75,975 | 88,265 | 87,620 | 86,143 | 80,733 | 94,619 |
| Total bp Deleted | 71,529,045 | 69,100,133 | 74,572,148 | 60,806,319 | 82,760,958 | 81,648,680 | 80,285,230 | 73,308,099 | 82,737,362 |

As with our finding for pangenome SVs, we found that linked-read deletions are most often shared between individuals of the same cave lineage rather than between cave lineages (Table 2), supporting current evidence that at least two cave lineages diverged from at least two separate surface lineages (Ornelas-García et al. 2008; Bradic et al. 2012; Herman et al. 2018; Moran et al. 2023). Thus, the number of deletions detected in each population is in line with our expectations based on the evolutionary history of *A. mexicanus*.

**Table 2.** Number of shared SV deletions between individuals from each pair of populations in linked-read sequencing dataset. Fewer deletions were shared between lineages than within lineages, supporting the best supported phylogenetic history of these populations. Deletions were considered shared if they had 90% reciprocal overlap between populations. This cutoff increases the confidence that the variants originate from a shared SV-generating event, rather than repeated origins in a similar location. Cells with no shading represent comparisons to surface populations, cells with mid-tone shading represent comparisons between cave lineages, and cells with dark shading represent comparisons within a cave lineage.

|  | Cave Lineage 1 |  |  | Cave Lineage 2 |  |  |  | Surface L1 | Surface L2 |
| --- | --- | --- | --- | --- | --- | --- | --- | --- | --- |
|  | Escondido | Molino | Vásquez | Pachón | Tigre | Tinaja | Japones | Jalpan | Rascón |
| Molino | 43,511 |  |  |  |  |  |  |  |  |
| Vásquez | 31,327 | 29,865 |  |  |  |  |  |  |  |
| Pachón | 24,301 | 23,098 | 24,905 |  |  |  |  |  |  |
| Tigre | 23,613 | 22,507 | 24,032 | 37,640 |  |  |  |  |  |
| Tinaja | 23,382 | 22,428 | 23,807 | 37,849 | 47,180 |  |  |  |  |
| Japones | 22,572 | 21,559 | 23,105 | 38,674 | 40,690 | 40,907 |  |  |  |
| Jalpan | 19,999 | 19,279 | 20,439 | 21,024 | 20,790 | 20,645 | 19,906 |  |  |
| Rascón | 24,917 | 23,904 | 25,407 | 31,565 | 32,211 | 32,341 | 30,424 | 22,512 |  |

### Identification of high-confidence cave-associated focal deletions

Deletions are likely contributors to cave-specific phenotypes given the loss of ancestral traits in cave-dwelling Mexican tetra and the high predicted functional impact of deletion variants according to SnpEff. Therefore, for downstream analysis we prioritized deletions over insertions. To further explore deletions segregating in Mexican tetra populations, we used the pangenome as a reference for the alignment and genotyping of 107 short-read WGS samples from nine total populations (Table S2). This approach allows SVs present in the pangenome to be confidently and accurately genotyped in resequenced samples using short reads; this would not be possible with traditional linear reference alignments (Hickey et al. 2020).

Our genotyping yielded 279,995 PGsr (pangenome + short-read WGS) deletions. To identify deletions statistically associated with cave-morph fish, we identified PGsr deletions present in each lineage and conducted lineage-specific runs of SnpSift CaseControl (Ruden et al. 2012). We used the allelic model and Fisher’s exact test with a Bonferroni-holmes correction to assess differences in deletion allele frequencies between cave populations (i.e., cases) and surface populations (i.e., controls). We identified 23,032 deletions in Lineage 1 and 6,606 deletions in Lineage 2 that were statistically associated with cave-morphs (adjusted p < 0.05; see methods for commentary regarding identification of fewer deletions in Lineage 2).

In line with best practices, we used multiple SV calling methods and retained SVs with consensus between callers to increase confidence in our PGsr cave-associated deletion calls (Ho et al. 2020). Of the deletions associated with cave-morphs by SnpSift CaseControl, 14,050 deletions in Lineage 1 and 5,246 deletions in Lineage 2 were supported by linked-read deletion calls.

Because we sought to explore deletions that have arisen since cave colonization and compare putatively independent deletion events between cave lineages, we isolated cave-specific deletions by removing deletions found in any surface populations (including all individuals from the linked-read and PGsr datasets). Finally, to identify deletions that have risen in frequency towards fixation, we required a deletion allele frequency of 0.8 or greater in at least one cave population. As a result, we obtained a final set of 2,431 unique deletions hereafter referred to as “focal deletions” (Table S3).

Notably, deletions identified as significantly associated with cave-morph fish in one lineage are not necessarily absent from the other; rather, they may exist at lower frequencies or lack sufficient genotype coverage to achieve statistical significance. Among the focal deletions in Lineage 1, 73% are unique to that lineage (i.e., not present at any frequency in Lineage 2), while 79% of focal deletions in Lineage 2 are unique (Table S4-A). Thus, approximately 26% of focal deletions are shared between lineages. These shared deletions may represent introgression between cave populations, reflect standing genetic variation that was undetected in our sampling of surface fish, or have arisen through similar *de novo* mutations.

In sum, we identify a set of 2,431 focal deletions that are strong candidates to contribute to cave-derived phenotypes. These deletions are significantly associated with cave-morph fish according to SnpSift, supported by both the PGsr dataset and linked-read dataset (i.e., are consistently identified across different SV calling methods), absent in all sampled surface fish, and high frequency (deletion allele frequency is ≥ 0.8) in one or more cave populations.

### Characteristics of focal deletions

Like the set of all deletions identified in the pangenome, the majority (96%) of focal deletions are ≤ 1 kb (mean = 202bp, median = 80bp; Table S4-B) and occur in CDS regions significantly less often than would be expected through random accumulation (only 14 focal deletions are within exons, permutation test, p < 0.003; Table S4-C). Again, this suggests that overarching purifying selection removes deletions that occur in coding regions. Focal deletions intersect introns and intergenic regions no more often than expected under random accumulation (permutation test, p = 0.303 and p = 0.376, respectively, Table S4-C). For focal deletions that impact protein coding genes, we do not find evidence that the distribution of the starting position of the deletion within the gene length differs from a uniform distribution (Chi-squared comparison, X = 9.098, df = 9, p-value = 0.4283; Table S4-D). Thus, focal deletions are not concentrated at certain positions along the gene length.

The position of deletions in the genome may be biased by the sequence context in that region. For example, repeated deletions within the pelvic enhancer region of stickleback fish are thought to be stimulated by stretches of TG sequence repeats which are highly mutagenic (Xie et al. 2019). Because TG repeats can promote deletion mutations, we explored whether any of our focal deletions occurred at TG repeat sites. We found six deletions in Lineage 1 and four deletions in Lineage 2 that contained a TG repeat region of 20bp or more (TG10+) (Table S4-E); however, none of the focal deletions at TG repeat sites are shared between lineages or occur within an overlapping region. We note one deletion of interest that occurs at a 54bp TG repeat (TG27), a 1,410bp deletion fixed in Molino and Escondido caves, that begins 5,413bp upstream of *ptprdb,* a gene associated with diabetes risk and insulin resistance in humans (Chang et al. 2012).

We identified only 13 shared focal deletions that are associated with the cave-morph in both lineages (i.e., only 13 focal deletions are significant by SnpSift in both lineages; Table S5-A) but find 67 instances of focal deletions in the same protein coding gene, but at different positions in different lineages (Table S5-B). Previous research has indicated that genes with large mutational target size are more likely to undergo repeated evolution (Gompel and Prud’homme 2009; Moran et al. 2023); correspondingly, we find that genes that repeatedly incurred unique deletions in each lineage are 3X longer (in terms of total bp) than the rest of genes that bear focal deletions (median length for genes with repeated deletions = 231,341bp, median length for all other genes impacted by focal deletions = 74,387bp, pairwise comparisons using Wilcoxon rank sum test with continuity correction p < 0.0001).

This further supports that longer genes are more subject to mutation and suggests that genomic biases such as length influence which genes are affected repeatedly in independent bouts of evolution.

### Putative functional impacts of focal deletions and association with cave-derived traits

We explored the potential functional impacts for genes containing focal deletions in both lineages through inspection of documented phenotypes in gene orthologs from zebrafish, humans, and mice (Table S5-C). Notably, we found that many genes containing deletions in both lineages have documented phenotypic impacts related to cave-derived phenotypes (Figure 5A).

**Figure 5.**
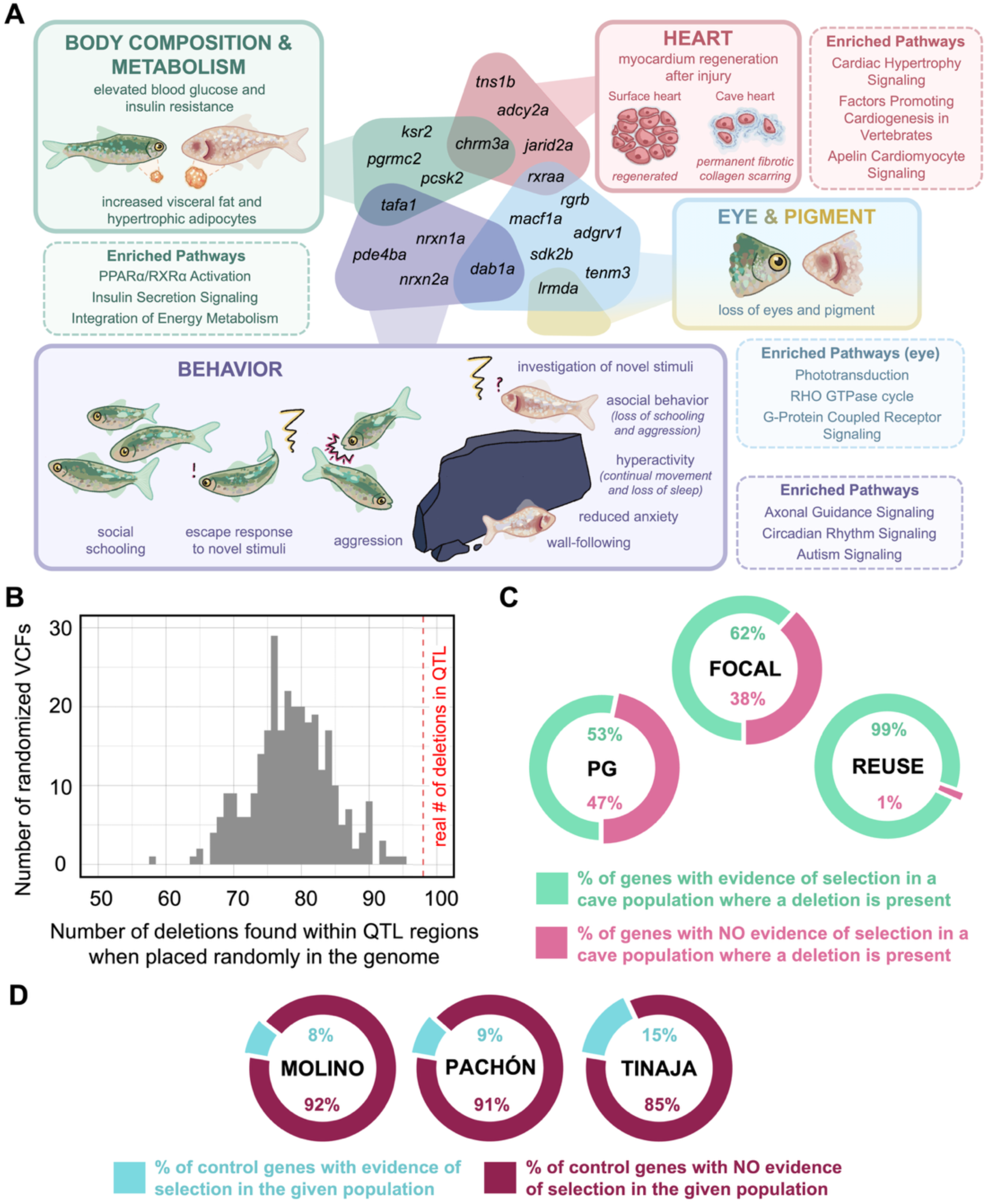
**A)** Putative relationships between genes containing deletions in both lineages and cave-derived phenotypes. Functional relevance for each gene was identified by exploring documented functional impact of the gene ortholog in zebrafish, mice, and humans, leveraging the databases Zfin, MGI, OMIM, and review of current literature (Table S5-C). Each colored bubble represents a suite of cavefish phenotypes, which are illustrated and compared to surface fish phenotypes in the corresponding boxes. The genes listed in each bubble have evidence of functional impact related to the cavefish phenotype indicated by the bubble. Examples of enriched pathways identified in Ingenuity Pathway Analysis of focal deletions pertinent to the phenotypes are given in dashed boxes (full list given on Table S7-A) **B)** Overlap between deletions and QTL from previous studies of the Pachón cavefish population (see Supplement S8-B). The histogram illustrates the number of deletions fixed in Pachón (Fst = 1 between Pachón and Rio Choy, n = 155 deletions) that were found within Pachón QTL regions in each run of a permutation test that produces a simulated VCF with deletions placed into the genome at random. The dotted line indicates the true number of of deletions within QTL regions (98 deletions). **C)** Proportion of genes under selection in three unique gene sets: PG = genes containing deletions that are present in the pangenome but did not have a statistical association with cave-morph fish (π = 16617), FOCAL = genes containing focal deletions that have a significant association with the cave-morph (n = 1260), REUSE = genes that contain a unique focal deletion in each cavefìsh lineage (n = 67). For a gene to be considered under selection, there must be evidence of a selective sweep in the gene in a cave population where the deletion is present. Genes are placed into the category based on their highest level of association (e.g., a gene containing both a PG deletion and a FOCAL deletion is put in the FOCAL category). **D)** Proportion of control genes (i.e., genes that contain no insertion or deletion SVs, n = 5265) under selection in each cave po

Additionally, we identified biological pathways overrepresented among the set of genes affected by focal deletions through Ingenuity Pathway Analysis (IPA) of human orthologs of Mexican tetra genes. We found 224 enriched pathways in Lineage 1 and six enriched pathways in Lineage 2 (Benjamini-Hochberg adjusted p-values < 0.05, Table S6-A). The smaller number of enriched pathways in Lineage 2 reflects the lower number of cave-associated deletions identified in this lineage resulting in fewer genes input for this analysis. Notably, all pathways enriched in Lineage 2 were also enriched in Lineage 1: RHO GTPase cycle, Autism Signaling Pathway, Synaptic adhesion-like molecules, ROBO SLIT Signaling Pathway, GABAergic Receptor Signaling Pathway (Enhanced), and Neurexins and Neuroligins. Despite this overall concordance in enrichment, mostly different genes in the same pathway incurred deletions between the two lineages (Table S6-B), indicating a pattern of convergence at the level of the pathway rather than convergence at the gene level. To consider cases of deletions affecting a gene’s regulatory sequence, we also ran pathway enrichment for genes with focal deletions within 5kb upstream (four enriched pathways in Lineage 1, one enriched pathway in Lineage 2, no shared pathways; Table S6-C) and 5kb downstream (no enriched pathways) of their coding sequence. Many enriched pathways correspond to the production of cavefish phenotypes including metabolism, eyes, heart, and behavior (when considering pathways significant in either lineage; Figure 5A).

Because most available RNAseq data are derived from Pachón cavefish and Río Choy surface fish, we investigated the impact of focal deletions on Pachón gene expression. WGS genotype data and RNAseq are not available for the same individuals, so we used only fixed focal deletions found in Pachón in cumulative exons and introns for the gene or within 5000 bp upstream and downstream of the gene in this analysis (obtained by filtering to only include deletions that have Fst=1 between Pachón and Río Choy surface). Correspondingly, for this test we restricted our set of control genes (i.e., genes that contain no SV insertions or deletions in the gene structure) to also exclude genes with SVs within 5000 bp upstream and downstream of the gene. We found that having a fixed focal deletion does not affect the likelihood that a gene is differentially expressed between Pachón cavefish and Río Choy surface fish in the eye at 54 h post-fertilization (Gore et al. 2018b) or liver tissues (Krishnan et al. 2020) compared to control genes (Fisher’s exact test, p > 0.05, Table S7-A). Because surface Mexican tetra can regenerate heart muscle after injury while cavefish retain permanent scarring, we also explored differential expression in heart tissues at 3, 7, and 14 days after injury (Stockdale et al. 2018). We found that genes with fixed focal deletions are more likely to be differentially expressed at 7 days after injury (Fisher’s exact test, p < 0.05) but not at 3 or 14 days after injury (Table S7-A). Additionally, we used a hypergeometric test that assesses whether a disproportionate number of differentially expressed genes contain fixed structural deletions in Pachón and obtain the same pattern of significance (p < 0.05 for hearts 7 days after injury, p > 0.05 for all other tissues; Table S7-A). We might have identified differential expression in deletion genes in only the heart samples because examining RNA-seq at multiple time points can better capture dynamic expression patterns that are missed in single time-point analyses like the liver and eye samples.

We identified 20 genes with fixed deletions that are differentially expressed in the eye, 30 in the liver, and 33 in the heart across all timepoints, as well as 14 genes with fixed deletions that are differentially expressed in multiple tissues (Table S7-A). Notably, in accordance with the eye loss phenotype seen in cavefish, 85% of differentially expressed deletion genes in the eye are downregulated in cave (Table S7-A). These results suggest that deletions do not ubiquitously affect expression, but a subset of deletions may have impacts that merit further examination.

Most QTL identified in Mexican tetra are derived from crosses between Pachón cavefish and Río Choy surface fish (Wiese et al. 2024). Therefore, we again used fixed focal deletions in Pachón to identify whether deletions were associated with regions of the genome thought to contribute to cave-derived traits. Using previously established QTLs in Pachón mapped to the surface reference by (Wiese et al. 2024), we ran a permutation test and found that fixed focal deletions in Pachón are found within Pachón QTLs more frequently than expected by chance, and significantly more frequently than expected based on the frequency of Pachón deletions in the pangenome (permutation test p < 0.005; Fisher’s exact test p < 0.0005; Table S7-B; Figure 5B). These results suggest that the focal deletions are enriched for genomic regions associated with cave-derived phenotypes, supporting their potential role in the genetic basis of these traits.

### Genes with cave-associated deletions are more likely to be under selection

To assess whether deletions associated with cave phenotypes are more likely to be under selection, we examined whether genes containing deletions also showed evidence of selective sweeps in the specific cave population(s) where the deletion is present. Evidence of selection was identified using diploS/HIC (Kern and Schrider 2018), a deep convolutional neural network approach that identifies sweeps from patterns of genetic variation. We compared three gene sets: PG genes (genes with deletions in the pangenome that are not statistically associated with cave-morph fish), focal genes (genes with deletions that are associated with cave-morphs, consistent with above usage of this term), and reuse genes (genes that contain focal deletions at different positions in each cavefish lineage). We found that the proportion of genes under selection increased significantly with the level of cave-association: PG genes had the lowest proportion under selection (53%), followed by focal genes (62%), and reuse genes had the highest proportion under selection (99%) (pairwise comparison of each gene set using Chi-squared test, all p < 0.0001; Table S5-D; Figure 5C).

Therefore, genes with cave-associated deletions are significantly more likely to be under population-specific selection than genes with deletions not associated with the cave-morph phenotype. Further, genes that harbor unique deletions in each cave lineage are more likely to have evidence of population-specific selection, supporting their potential role in the repeated evolution of cave-adapted traits.

To provide an additional comparison, we explored how often control genes with no insertion or deletion SVs had evidence of selection in each population. Only 8%, 9%, and 15% of control genes had evidence of selection in Molino, Pachón, and Tinaja, respectively (Figure 5D; Table S5-D), substantially less than the proportion of genes with deletions under selection. This result further supports the hypothesis that structural variation, particularly deletions, contributes to adaptation in cavefish.

The Mexican cavefish system, characterized by extreme environmental pressures and repeated evolution of cave-morph fish, offers a uniquely powerful framework for uncovering the evolutionary impact of SVs. By combining pangenome driven SV detection with population-specific selective sweep scans, our study reveals that not only do structural deletions likely contribute to the production of cavefish phenotypes, but they do so through repeated natural selection.

## DISCUSSION

Thus far, comprehending the genetic basis of cave adaptation, driven by the strong selective pressures of darkness, food scarcity, and low oxygen, has involved examinations of single nucleotide variation. Our understanding of the impact of structural variation on cave adaptation was concentrated on the unique deletions in the *oca2* gene between cavefish lineages that play a dual role in both depigmentation and loss of sleep (Protas et al. 2006; Bilandzija et al. 2013; O’Gorman et al. 2021). In this study, we generated a pangenome reference for the Mexican tetra, *Astyanax mexicanus*, representing one of the first pangenome graphs for naturally evolving populations (but see, Ruggieri et al. 2022; Secomandi et al. 2023; Fang and Edwards 2024; Edwards et al. 2025; Quah et al. 2025). Our pangenome approach enabled us to comprehensively characterize structural variation between and within independently evolved lineages. Through our pangenome-guided investigation of deletions, we identified a high-confidence set of 2,431 focal deletions associated with cave-morph fish. These deletions are consistently identified across different SV calling methods, absent in all sampled surface fish, and high frequency (deletion allele frequency is ≥ 0.8) in one or more cave populations.

Our investigation of deletions in Mexican tetra revealed a remarkable pattern of unique focal deletions within the same gene between cavefish lineages. There often exist multiple mutational routes to a phenotypic outcome, and this genotypic redundancy results in a reduced likelihood that the same gene should be repeatedly used in independent bouts of evolution (Yeaman et al. 2018). It is therefore especially interesting to find the same gene altered repeatedly, suggesting constraints guiding evolutionary trajectories, and/or particularly advantageous fitness outcomes of changes at certain loci (Storz 2016). Beginning with Darwin, the evolution of loss-of-function traits in cave organisms has historically been attributed to drift and disuse rather than selection for loss (Darwin 1859; Wilkens 1988; Leys et al. 2005). However, genomic analyses of Mexican cavefish have shown that selection, as well as drift and relaxed constraint prominently influence the evolution of cave-derived traits (Moran et al. 2023; Roback et al. 2025). Here, we found that while only 8%, 9%, and 15% of control genes with no SVs had evidence of selection in Molino, Pachón, and Tinaja, respectively, 99% of genes that repeatedly incurred unique deletions in each cave lineage had evidence of selection in the populations where the deletion is present. These findings suggest that the presence of unique high-frequency deletions in the same gene in each lineage is not due to chance or relaxed constraint simply allowing these variants to persist, but repeated selection favoring independent mutations in these particular genes in caves.

As further evidence that genes where deletions have occurred and risen in frequency repeatedly may play an important role in the production of cavefish traits, we found that genes with repeated deletions in each lineage have documented phenotypic impacts in model systems related to many cave-derived phenotypes (see Table S5-C). These traits include those related to the physiological and behavioral resilience to nutrient limitation.

Given the lack of light, and therefore photosynthetic primary production, caves are inherently nutrient limited environments. External nutrient input may arise through bats living in the caves or seasonal flooding (Culver and Pipan 2019), however, it is likely that many caves containing Mexican cavefish are consistently nutrient limited (Wilson et al. 2021). Correspondingly, cavefish can maintain body weight when starved and exhibit hyperphagic behavior when food is present (Aspiras et al. 2015). They also have enlarged fat cells and greater visceral fat (Xiong et al. 2018; Xiong et al. 2022), as well as insulin resistance and elevated blood glucose (Riddle et al. 2018). We highlight two genes with repeated deletions between lineages and evidence of selection, *pcsk2* and *ksr2,* with convincing connections to these phenotypes. *Pcsk2* has an 81bp deletion in the fifth intron (of 11 total introns) in Lineage 2, followed by an 59bp deletion in the last intron in Lineage 1. Reduced levels of *pc2,* the mammalian ortholog of *pcsk2*, is implicated in Prader-Willi syndrome in humans, which is a disease characterized especially by insatiable hunger and often abnormal eyesight (Butler 1990; Gabreëls et al. 1998; Bohonowych et al. 2021). Additionally, in *pc2* null mice processing of insulin and glucagon is impaired (Furuta et al. 1997; Furuta et al. 1998; Furuta et al. 2001). *Ksr2* contains three focal deletions, a 67bp deletion in Lineage 2 in the first intron (of 22 total introns), a 57bp deletion in Lineage 1 in the sixth intron, and another 66bp deletion in Lineage 2 in the penultimate intron. Human *ksr2* is located within a region associated with both obesity and type 2 diabetes (Bowden et al. 1997; Li et al. 2004). In mice, *ksr2* knockouts are obese, glucose intolerant, and exhibit hyperphagia that is unresponsive to leptin, a hormone that signals feelings of fullness and consequently reduces hunger (Revelli et al. 2011). In fact, *ksr2* knockout mice are significantly more obese and glucose intolerant than *mc4r* knockout mice, a gene with previously established causative SNP in cavefish (Revelli et al. 2011; Aspiras et al. 2015).

One phenomenon likely shaping our observed pattern of gene reuse is the presence of repeated environmental pressures. Repeatedly altered genes in multiple evolutionary events are also seen in the evolution of pelvic spine loss in freshwater threespine stickleback (Chan et al. 2010), adaptation of Poeciliidae fish living in springs rich in toxic hydrogen sulfide (Greenway et al. 2020), and in waterfowl adapted to high-altitude hypoxia (McCracken et al. 2009), all systems with environmental pressures that are similar in each bout of adaptation. Results from experimental evolution have also demonstrated parallel genomic evolution in populations evolved in the same environment (Bailey et al. 2015), further suggesting that repeated environmental pressures are likely key influences on gene reuse.

In addition to shared environmental pressure, characteristics of the genomic sequence may introduce biases that predispose certain genes for reuse. In line with previous research indicating that genes with large mutational target size are more likely to be subject to repeated evolution (Gompel and Prud’homme 2009; Moran et al. 2023), we find that genes that have repeatedly undergone unique deletions in each lineage are 3X longer than the rest of genes in our set of focal deletions. This result further supports that longer genes offer more opportunity for mutations and suggests that genomic biases, such as gene length, influence tendencies for genes to be affected in repeated bouts of evolution.

When considering all focal deletions, not just those in the same gene between lineages, we find that fixed focal deletions in Pachón occur non-randomly in regions of the genome within established QTLs for cave-derived phenotypes, supporting that these variants may contribute to the genetic basis of cavefish traits. Additionally, multiple biological pathways pertinent to cavefish traits are overrepresented among the set of genes affected by focal deletions. Among these pathways are Circadian Rhythm Signaling, Melatonin Signaling, Insulin Secretion Signaling, Type II Diabetes Mellitus Signaling, Phototransduction, G-Protein Coupled Receptor Signaling, and Cardiac Hypertrophy Signaling. These pathways have clear connections to cavefish traits of circadian rhythm disruption and sleep loss, insulin metabolism, eye loss, and heart regeneration loss. Interestingly, we find that for pathways enriched in both lineages, deletions mostly affected different genes in the pathway in each lineage. This result illustrates that there also exists genetic convergence at the level of the pathway, but not at the precise level of the gene, in the repeated evolution seen in this system.

Among the genes harboring focal deletions, we identified a set of 14 genes that are differentially expressed between cave and surface in multiple tissues, suggesting that they may have pleiotropic effects, impacting multiple cave phenotypes at once. One gene, *ndst1b*, contains a deletion in Lineage 2 caves, is downregulated in cavefish in eyes, liver, and heart, and is associated with QTLs in Pachón for eye size, dentition, and body condition (Protas et al. 2007; Protas et al. 2008). In zebrafish and mice, knockdown of *ndst1b* causes altered neural crest development, with consequences for craniofacial development, eye defects, shortened body and pectoral fin length, and alterations to chromatophore distribution (Grobe et al. 2005; Filipek-Górniok et al. 2015), consistent with the changes in craniofacial and fin morphology and pigment loss in Mexican cavefish (Yamamoto et al. 2003; Protas et al. 2008; Powers et al. 2017; Powers et al. 2018; Atukorala et al. 2019). Neural crest cells play a key role in the development of multiple cave phenotypes, and transplantation of cranial neural crest cells from surface fish results in increased pigmentation and eye size in cavefish (Yoshizawa et al. 2018). The deletion identified in *ndst1b* occurs in the second intron, and *ndst1b* is significantly downregulated in multiple tissues in cavefish, suggesting that the deletion may have a regulatory effect and could be relevant to multiple cavefish phenotypes. This deletion, and many others (including those discussed prior, highlighted in Figure 5A, and on Table S5) represent exciting targets for further research using transgenesis and gene-editing technology (Stahl et al. 2019) to better understand the functional contribution of deletions to cavefish phenotypes.

In this study, we demonstrate that SVs are non-randomly distributed across the genome and provide further evidence that longer genes represent larger mutational targets for SVs and are more often subjected to recurrent disruption during independent evolutionary events. We identify patterns of convergence at the level of the pathway and the gene and provide multiple lines of evidence that deletions may contribute to the emergence of cave-derived phenotypes including that: 1) fixed deletions occur non-randomly within established QTL regions associated with cave traits, 2) focal deletions affect genes with known roles in cave-like traits, and 3) focal deletions are enriched in relevant biological pathways. Furthermore, genes harboring deletions are more likely to bear signatures of selection compared to control genes without SVs. Strikingly, while 8-15% of control genes had evidence of selection in cave populations, 99% of genes that have repeatedly incurred unique deletions across cave lineages show evidence of selection in caves, suggesting that this pattern is not simply the result of relaxed constraint, but reflects repeated, lineage-specific selection on parallel genetic changes. In sum, our results support a model in which deletions may be contributors to adaptive trait evolution, enabling convergent solutions to similar ecological challenges.

## METHODS

### Pangenome construction and processing

We built a pangenome from autosomal sequences of the surface and three cave morph *Astyanax mexicanus* assemblies AstMex3_surface (Río Choy), AstMex3_Molino (Molino cave), AstMex3_Tinaja (Tinaja cave), AMEX_1.1 (Pachon cave) (Imarazene et al. 2021; Warren et al. 2024). We ran the minigraph-cactus pipeline (Hickey et al. 2024) using the cactus v2.4.2 Docker image as described in (Rice et al. 2023) (see Data Availability Section). We specified the Río Choy surface assembly as the reference, due to it being of the highest quality as assessed by assembly representation and contiguity metrics (Warren et al. 2024). The graph was output in multiple formats including GFAv1.1, hal, vg, and vcf.

As default in cactus 2.4.2, the vcf was post-processed with vcfbub which exposes the snarl tree to annotate top-level (LV=0) and nested variants (LVs > 0). We subset the graph to only include top-level variants (‘vcfbub -l 0’) for all subsequent analyses. Although an updated version of Minigraph-Cactus left aligns this vcf with ‘bcftools norm’, the version used in this study (v2.4.2) lacked normalization as a default. We initially post-processed the top-level vcf with ‘bcftools norm’, but in doing so observed negative impacts of the left alignment, wherein variants were reassigned new coordinates that made them both incompatible with other formats of the graph (e.g. hal, gfa, vg), and incompatible with one another (e.g. an individual cannot have two different alternate genotypes at the same site; https://github.com/ComparativeGenomicsToolkit/cactus/issues/1557). We chose to use the top-level vcf without left alignment for all subsequent analyses. As a final post-processing step, we subset the top-level, unnormalized vcf to lines where at least one individual (Molino, Pachón, or Tinaja) had an alternate genotype. Given that our input assemblies are haploid compressed genome assemblies, each ‘genotype’ in the pangenome is a single allele (e.g. ‘1’ instead of ‘1/1’, for example).

### Pangenome genotyping

In the post-processed vcf, variant types were categorized based on the length of the alternate (ALT) allele and the length of the reference allele (REF) after splitting multiallelic sites (Data Availability Section). Structural variants were defined as those greater than or equal to 50bp and up to 100kbp, in accordance with other pangenome approaches e.g. (Fang and Edwards 2024). To set equal thresholds across insertions and deletions, we categorized the structural insertions >100kbp as “large structural insertions”. Similarly, 1bp indels, small insertions (2bp – 49bp), and small deletions (2bp - 49bp), were defined through inspection and subtraction of the ALT and REF alleles. We defined 1bp indels as 1 base pair sequence gains or sequence losses that do not, otherwise, meet the criteria of being a multi-nucleotide polymorphism. The code for parsing and the thresholds used for other variant type categories can be found in the Data Availability Section.

### Pangenome summary statistics

Variants on unplaced scaffolds were excluded from pangenome summary statistics reported in the paper. Throughout the paper, gene is defined in accordance with NCBI definition used for GTF files, where ‘gene’ describes coordinates encompassing all isoforms of a gene (the union of all introns and exons). For this study, we only analyzed protein coding genes. Statistics regarding the relationship of structural variants to different genomic features, other than repetitive elements, were calculated based on the annotation of the pangenome vcf (see *Annotating genomic features of structural variants*). Calculations of gene length, proportion of gene deleted, and position of deletions within a gene were all calculated based on the gene coordinates specified in the GTF.

### Identification of repetitive regions within insertions

We used RepeatModeler 2.0.4 (Flynn et al. 2020) with the -LTRStruct option to create libraries of transposable elements for the surface and each of the cave morph genome assemblies. For each assembled cave morph, we subset the VCF to contain only insertions present in that genome assembly. We converted column 5 of the resulting VCF to a fasta file and used this, along with the repeat-families.fa output of RepeatModeler as described above, as input to RepeatMasker v4.1.7-p1 (Tarailo-Graovac and Chen 2009) with default parameters.

### Annotating genomic features of structural variants

To identify the impacts of structural variants in *A. mexicanus*, we annotated the pangenome vcf using SnpEff 5.2a (Cingolani et al. 2012) with informed breakpoints. The SnpEff database was built from the AstMex3_surface reference using the NCBI GTF. Custom annotations were provided for population-specific sweeps, and QTLs using the –interval option. Additionally, we added the tag ‘CDSID:’ using coordinates from AstMex3_surface NCBI GTF with the –interval option. We used QTL coordinates from (Wiese et al. 2024). QTL annotations were added using the tag “[population]QTLs:” followed by the associated trait. After generating coordinates for cave-specific selective sweeps using DiploS/HIC (see Methods: *Detection of genome-wide selective sweeps*), we annotated the vcf for sweeps using the tag “[Population]Sweep:selection.present, [Population]Sweep:selection.absent, or [Population]Sweep:NA”.

### Short-read WGS genotyping of surface and cave populations

We mapped and genotyped short-read surface and cave morph population data (full information and sequence accessions available on Table S2) to the pangenome graph described in this study using vg giraffe with default options (Hickey et al. 2020). GAM format files were surjected to BAM format with surface serving as the reference genome using the command “vg surject” with default options. Only structural variants within 50bp - 100kbp were retained for downstream analyses. The location of our genotyping scripts and postprocessing, along with all other analyses described in this paper, can be found in the Data Availability section.

### Linked reads generation and SV genotyping

For validation of PGsr deletions, we used 10X Genomics linked-read sequencing of nine individuals from two surface populations (Jalpan and Rascón) and seven caves (Escondido, Molino, Japones, Pachón, Tigre, Tinaja, and Vásquez). All processing of the linked-read data and SV calling was completed using Long Ranger (v2.2.2) (Zheng et al. 2016). We generated FASTQ files from the Illumina binary base call sequence output by running longranger mkfastq and created a 10X compatible reference from the *A. mexicanus* surface fish reference genome (GCF_023375975.1) using the longranger mkref command. We then ran Long Ranger in whole genome mode (wgs). The longranger wgs command performs alignment of sequences to the reference genome, de-duplication and filtering, and uses the Chromium molecular barcodes to call and phase SNPs, indels, and structural variants. From the phased structural variants called by Long Ranger we retained only variants 50bp-100kb in length that passed all filters tested in the pipeline. These filters use barcode and phasing information to improve the quality of variant calls and reduce false positives (description of filters in the Long Ranger pipeline: https://support.10xgenomics.com/genome-exome/software/pipelines/latest/output/vcf). To determine the number of shared deletions between each population pair we used bedtools intersect (Quinlan 2014) to identify and count deletion SV calls with ≥90% reciprocal overlap. This level of overlap increases the likelihood that deletions being counted as shared are indeed comparable variants rather than unique deletions in similar positions.

### Identifying a high confidence set of cave candidate deletions

We integrated short-read genotyping with linked-read genotyping, employing stringent filtering criteria and a statistical test to identify SVs associated with cave-morph fish. First, we identified deletions in the PGsr dataset statistically associated with cave-morph fish using SnpSift (Ruden et al. 2012). The allelic model implemented in SnpSift’s CaseControl analysis (v5.2) utilizes Fisher’s exact test to assess statistically significant associations between genotype and phenotype. This test is effective in identifying variants exhibiting a high frequency of alternate alleles in case samples with a high frequency of reference alleles in control samples. We applied this test in a phylogenetically informed framework, treating cave samples as the ‘case’ and surface samples as the ‘control.’ To account for the independent origins of the cave phenotype in Lineage 1 and Lineage 2, we performed separate analyses for each lineage using the post-processed PGsr data. Only SV deletions were included as input, with multiallelic sites joined. We adjusted p-values with Holm-Bonferroni correction using a custom code written with pysam (v. 0.22.1; https://github.com/pysam-developers/pysam) and the multipletests() function in statsmodels 0.14.0 (Seabold and Perktold 2010), setting the significance threshold to adjusted p = 0.05. The code to correct p-values from SnpSift can be found in the Data Availability Section.

We likely identified fewer significant deletions in the Lineage 2 SnpSift CaseControl analysis due to 1) reduced statistical power resulting from fewer surface individuals in Lineage 2 (Lineage 1 = 19 controls, 31 cases; Lineage 2 = 12 controls, 45 cases); and 2) a reference bias wherein a multiallelic site with alternate Rascón allele (the L2 surface population) would be excluded by SnpSift due to the absence of reference genotype in the control samples. We also note that variants deemed significant in the SnpSift CaseControl analysis are only those for which there was high genotype coverage resulting from mapping the short-read WGS to the pangenome. Therefore, we can confidently say these deletions are associated with cave-morph fish, however, there are likely additional deletions that are potentially biologically relevant, but do not meet our threshold for significance. For instance, deletions in *oca2* are well documented in cavefish (first identified in (Protas et al. 2006), and present in our PGsr deletion dataset, but are not identified as significant through SnpSift CaseControl due to reduced genotype coverage in our samples.

In investigations of structural variation, the confidence of SV calls is increased using multiple SV calling methods and retaining calls with consensus between programs (Ho et al. 2020). Generally, a 50% reciprocal overlap between callers is used to determine high-confidence SV calls (Sudmant et al. 2015; Audano et al. 2019; Kosugi et al. 2019; Ho et al. 2020). We generated a set of high-confidence SV deletions from PGsr genotyping by requiring a ≥50% reciprocal overlap with a deletion call from Long Ranger in a linked-read cave sample from the same lineage. To further isolate cave-specific deletions, we employed bedtools intersect to filter out SV deletions in cave samples that showed ≥50% reciprocal overlap with SV deletions in the linked-read surface samples (Jalpan for Lineage 1 and Rascón for Lineage 2).

We calculated the population-specific frequency of our high-confidence deletions using a custom perl script (see Data Availability Section). We then filtered our set of high-confidence deletions, requiring an alternate allele frequency of 0 in all sampled surface populations (Lineage 1: Río Choy, Mante, and Lineage 2: Rascón). The final sets of Lineage 1 and Lineage 2 cave-associated SV deletions were then annotated with SnpEff using the same approach applied to the pangenome.

### Detection of genome-wide selective sweeps

To find genomic regions possibly associated with adaptation, we identified regions of the genome with evidence of selective sweeps using diploS/HIC (Kern and Schrider 2018). This method combines a convoluted neural network approach and calculation of population genetic statistics from sequence data to infer selection. DiploS/HIC requires empirical population genetic information in VCF format. We generated this input using srWGS data from two surface populations (Río Choy n=9 and Rascón n=13) and three cave populations (Molino n=14, Pachón n=19, and Tinaja n=21) (sequence accessions available on Table S2-B). We first performed genotyping against the Río Choy surface reference (Warren et al. 2024) using Genome Analysis Tool Kit (GATK) (Van der Auwera and O’Connor 2020) v4.4.0 GenotypeGVCFs tool to produce a VCF file for each population. We then subset these VCFs to only include SNPs and added an additional mask to remove any SNPs that occurred within indels or a 3bp buffer region on either side. We followed GATK best practices (https://gatk.broadinstitute.org/hc/en-us/articles/360035890471-Hard-filtering-germline-short-variants) to hard filter SNPs and removed SNPs that occurred within repetitive regions using WindowMasker files downloaded from NCBI.

Next, we simulated population genetic data under neutral, soft, and hard sweeps in discoal (Kern and Schrider 2016), using the same demographic parameters as in a past run of diploS/HIC in Mexican tetra (Moran et al. 2023). We created separate simulated datasets for surface and cave to account for past population size changes and reduce the potential for false positives resulting from population bottlenecks that can cause neutrally evolving genomic regions to exhibit low genetic variation mimicking a selective sweep. We then generated summary statistics and feature vectors for the simulated surface and cave datasets which we used to train and test the convoluted neural network and produce a model for cave and surface. Finally, we supplied our VCFs containing real population genetic data to the trained diploS/HIC model which generated another set of summary statistics and feature vectors which were used to predict selection in 5kb windows across the genome. Each window was classified as being neutral, containing hard sweeps or soft sweeps, or being linked to a region that experienced a hard or soft sweep. Although diploS/HIC is a powerful approach to detect regions of the genome containing selective sweeps versus those that do not, it is not as reliable at distinguishing hard sweeps from soft sweeps (Kern and Schrider 2018). For this reason, we grouped hard and soft sweeps as evidence of selection and do not distinguish between the two. To further increase the confidence of our selection calls, we then swapped the underlying demographic models (running the cave population genetic dataset with the surface demographic model and vice versa) and retained only windows with consistent calls between models.

### Permutation testing of SV prevalence

To assess how the distribution of structural deletions across different genomic features (CDSs, introns and intergenic regions) differ from chance, we ran a permutation test using a custom script. This test randomizes positions in the VCF, restricted to the lengths of the chromosomes and generates a user-specified number of permuted VCFs (Data Availability Section). We generated 300 permutations of pangenome structural deletions. We annotated these with SnpEff following the methods previously described (see *Annotating genomic features of structural variants*). We also ran permutation tests on Lineage 1 and Lineage 2 focal deletions to test relationships of cave-associated deletions to selective sweeps in cave populations, QTLs, CDSs, introns and intergenic regions. We used a Fisher’s exact test to ascertain how the distribution of focal deletions differs from overall population-specific deletions in the pangenome.

### Orthology inference

We used Orthofinder (v3.0.1b1; Emms and Kelly 2019) to identify human orthologs of genes with focal deletions. Briefly, we used the same methods and species sampling as our prior study (Warren et al. 2024) but, whereas the prior study used Ensemble gene predictions as input, here we used NCBI gene predictions. The metadata associated with input species / annotations, and a summary of orthogroup assignments is reported in Table S8. Assignment of gene symbols to corresponding stable identifiers was accomplished with scripts reported in our prior study (Warren et al. 2024).

### Pathway analysis

We performed signaling pathway overrepresentation tests for statistical significance using Ingenuity Pathway Analysis (IPA) (Krämer et al. 2014). For each lineage, we identified three sets of genes: 1) genes with deletions within an intron or exon, 2) genes with deletions within 5kb upstream of the start codon, and 3) genes with deletions within 5kb downstream of the stop codon. For each Mexican tetra gene, we identified the human ortholog from our Orthofinder analysis. Genes with single copy human orthologs were used as input for IPA because *A. mexicanus* is unavailable in this software. Each of the three gene sets was run independently for each lineage (six runs in total). Significant pathways were those that had a Benjamini-Hochberg adjusted p-value < 0.05. To not inflate enrichment, we restricted pathway enrichment analysis to single copy human orthologs (including 1:1 and multi:1 orthologs between *A. mexicanus* and *H. sapiens*). This is consistent with methods in functional annotations overall, which rely on orthology relationships beyond 1:1 orthology (e.g. (Hernández-Plaza et al. 2022). We note that because multiple Mexican tetra genes may map to the same human ortholog, the number of genes within a pathway that appear to be impacted in both cavefish lineages is overrepresented (i.e., different cavefish genes with deletions may map to the same human ortholog, which can give the false impression that the same gene is affected in both lineages).

### Quantification of cave vs surface gene expression

Previously published RNAseq datasets were utilized for differential gene expression: wild-type livers (Krishnan et al. 2020), post-injury hearts across three time points (Stockdale et al. 2018), and 54 hours post fertilization eyes (hpf) (Gore et al. 2018a). Reads were mapped using the Amex 3.0 NCBI annotation with Star v2.7.11b (Dobin and Gingeras 2015). We enabled –quantmode to count reads during mapping, which generates identical results to HTseq (Anders et al. 2014). We did not trim reads given that trimming is redundant when mapping involves soft-clipping (Liao and Shi 2020), a default in the STAR aligner. We used DESeq2 to normalize raw counts based on size factors, remove outliers based on Cooke’s distance, perform differential expression, and run PCAs on variance stabilized counts (Love et al. 2014). All comparisons were performed with Pachón vs Río Choy, as both populations were available in all datasets. For the livers, eyes and hearts the significance threshold was set to adjusted p-value less than 0.05.

### Associating candidate deletions with gene expression

Genotyping of the individuals used for RNAseq is not available. Therefore, to ensure the deletion was present in the Pachón individual used for RNAseq, we calculated Weir and Cockerham’s Fst using vcftools on the Lineage 2 focal deletions (Danecek et al. 2011). We filtered the dataset to only include variants in Pachón that have Fst=1 compared to Río Choy. We then identified all genes with a fixed focal deletion that intersects the gene structure, 5000 bp upstream or downstream from the start and stop site. In addition, we identified a set of control genes that did not have any evidence of structural variation (including both insertions and deletions) within the gene structure, or 5000 bp upstream or downstream from the start and stop site.

We first tested whether the likelihood that a gene is differentially expressed differs between fixed focal deletions and control deletions in each tissue using a Fisher’s exact test. Additionally, we tested whether fixed focal genes are differentially expressed more frequently than expected by chance in each tissue using a hypergeometric test with the phyper() function in R, setting lower.tail=FALSE. This calculates whether candidate genes are overrepresented amongst the genes that are differentially expressed between cave and surface. Differentially expressed genes that are focal genes are treated as ‘successes’ in the test. The significance of the rate of success is based on N= the number of protein coding genes in the genome, K= the number of differentially expressed genes, M= the number of candidate genes, and x= the number of differentially expressed genes that are focal genes (i.e. ‘successes’).

## ACKNOWLEDGEMENTS

The High-Performance Computing systems of the Minnesota Supercomputing Institute (University of Minnesota) and Hellbender (University of Missouri) enabled this computationally intensive research.

## AUTHOR CONTRIBUTIONS

**E.Y.R.** led the project, including study design, data analysis, manuscript writing and editing, and completed all figure design. **M.X.** contributed to study design, data analysis, and manuscript preparation. **E.S.R.** contributed to study design, data analysis, and manuscript review. **A.W.** performed the IPA analysis. **R.A.C.** provided technical support and manuscript review. **C.G.E.** contributed to manuscript review. **A.C.K. and N.R.** contributed to project ideation and manuscript review. **S.E.M. and W.C.W.** initiated project ideation, supervised the project, and contributed to study design, analyses, manuscript writing and editing.

## DATA AVAILABILITY

The reference genomes used in this study can be found through NCBI at the following accessions: GCA_023375975.1, GCA_023375835.1, GCA_019721115.1, GCA_023375845.1.

All other data, including sequence accessions for WGS and 10X samples, custom scripts, and analysis pipelines can be found within the Supplementary Materials and at the following repositories: https://github.com/WarrenLab/minigraph-cactus-nf https://github.com/robackem/Cavefish-SV https://github.com/maggs-x/Amex-pangenome

---

**Supplemental Figure 1.**
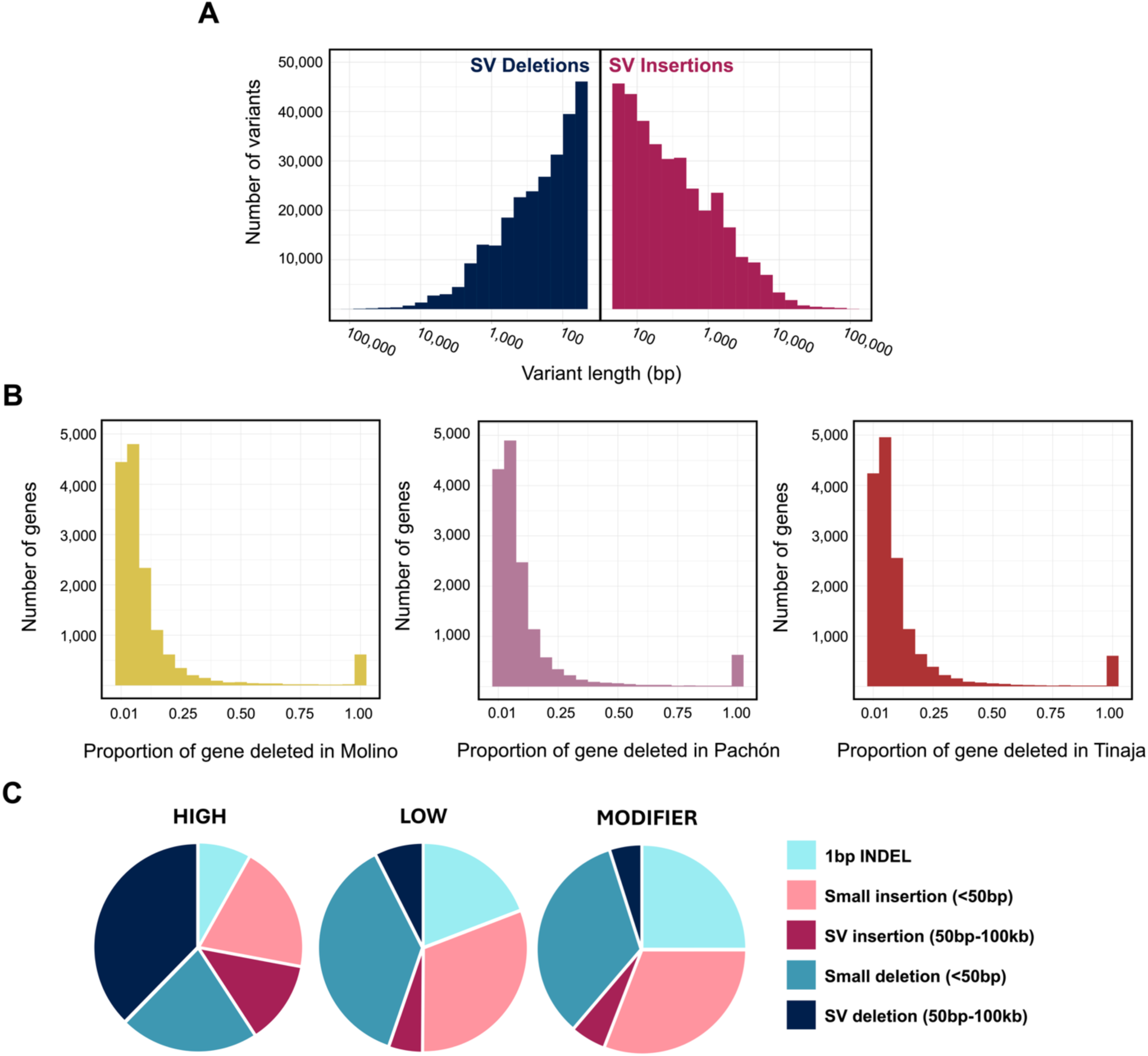
Further information on panqenome insertions and deletions. **A)** Length of SV deletions (left) and insertions (right) found in the pangenome. Variant length represented on a loglũ scale. B) Proportion of gene deleted. For protein coding genes impacted by deletions, the proportion of the gene structure (including introns) that falls within the deletion is given for each population (Molino, Pachón, Tinaja). If a gene contained more than one deletion in a population, the number of bp deleted was added for all deletions to generate a value of bp deleted across the gene in total. C) Predicted impact of pangenome insertions and deletions from SnpEff annotation. Variants with HIGH predicted impacts include partial or whole exon deletions, variants causing frame shifts (i.e., when indel size is not multiple of 3), alteration of splicing donor or acceptor region, stop codon gain or loss, start codon loss. Variants with LOW predicted impacts include production of upstream start codon in 5’UTR, synonymous codon changes including synonymous start or stop, alterations of splice region (outside of donor or acceptor region). Variants with MODIFIER predicted impacts include variants within 5kb upstream or downstream of a gene, variants within 5’ or 3’UTRs, and variants in intergenic regions or introns.

